# First report of *Photobacterium damselae* subspecies *damselae* in Razorbill (*Alca torda*)

**DOI:** 10.1101/2025.05.15.654261

**Authors:** Adriano Minichino, Francesca Lucibelli, Tullia Guardia, Rosario Balestrieri, Serena Aceto, Emanuela Vaccaro, Ludovico Dipineto, Marzia Sapio, Antonio Santaniello, Luigi Maria De Luca Bossa, Nicola D’Alessio, Alessandro Fioretti, Luca Borrelli

## Abstract

A razorbill (*Alca torda*) was found dead in Bacoli, Italy, on January 16, 2023, during an exceptional irruptive event. The bird, a young female weighing 800 g with a moderate nutritional status and no external traumatic injuries underwent a post-mortem examination, which revealed coelomitis with severe congestion of the liver, lungs, kidneys, and myocardium. Lung, liver, kidney, and heart samples were collected for microbiological and histopathological analyses. Bacterial isolation on blood agar showed the growth of spherical/ovoid colonies after 48 hours, consistent with the *Photobacterium* genus. Identification by MALDI-TOF MS and PCR confirmed the presence of *Photobacterium damselae* subsp. *damselae* (Pdd), supported by 16S gene sequencing and detection of the *ureC* gene. PCR screening for virulence factors identified the *hlyAch* gene in lung samples, suggesting a potentially pathogenic strain in avian species. Histopathological examination showed severe, inflammatory infiltrates and widespread hemorrhages in the liver, the lung tissue and kidneys. In the myocardium, mild and multifocal lymphocytic infiltrates were present. These findings suggest a significant role of *P. damselae* subsp. *damselae* in the observed lesions. The results of the study revealed a new cross-species transmission of this potential zoonotic bacterium and the need for further research in this field.

**Importance:** With this study we revealed the first isolation of *Photobacterium damselae* subsp. *damselae* in a razorbill but especially the first in a migratory bird. We documented a new cross-species transmission of *P. damselae* subsp. *damselae* and the need for further research, highlighting its zoonotic potential and therefore raising concerns about potential implications for marine wildlife, aquaculture and human health. Considering the role of migratory birds in the spread of infectious diseases even over long distances and the effect of climate change on marine ecosystems, and consequently on the circulation of pathogens, it is essential to adopt a preventive approach to mitigate this potential zoonotic risks.

## Introduction

The razorbill (*Alca torda*, Linnaeus 1758) is an auk species with breeding colonies located in the boreal and sub-Arctic waters of the North Atlantic Ocean. This species primarily inhabits shallow coastal waters. Globally, the razorbill population is estimated at approximately 700,000 breeding pairs, with the majority (around 80%) concentrated in Iceland and the British Isles (12). Following the breeding season, razorbills disperse to open waters of the North Atlantic where they typically overwinter (14). Additionally, southward movements have been documented, with individuals redistributing across the Atlantic coasts of France and Portugal (20). These movements can extend as far south as the Canary Islands and the western Mediterranean Sea, with razorbills occurring in small numbers (6). Occasionally, irruptive events may occur, resulting in unusually high numbers of specimens entering the Mediterranean Sea and dispersing across the coastlines of countries like Spain, France, Italy, Algeria, Tunisia, Libya, Malta and Greece (2, 5). It has been suggested that clupeids, such as the capelin *Mallotus villosus*, the Atlantic herring *Clupea harengus* and sandeels (*Ammodytes* spp.), due to their relevant content of fat, represent an important energy source for razorbills, especially during winter (3, 14). During the last mass irruption into the Mediterranean Sea in 2022/2023 the razorbills adapted to the local fish prey by targeting species of similar shape and size as those that make up the core of their diet in their main wintering areas of the Atlantic. Moreover, they showed peculiar feeding behaviours such as foraging close to shorelines or even within ports, as documented by photos and videos made by citizens (2, 16).

## Materials and Methods

On January 16, 2023, a razorbill was found dead and beached in Bacoli (Naples), Italy. It was subjected to a post-mortem examination. The razorbill weighed 800 grams and was identified as a young female with a moderate nutritional status. No evidence of trauma was observed during the external examination. Regarding the examination of the coelomic cavity, coelomitis was found with severe congestion of the liver, lungs, kidneys and myocardium. Samples of the liver, lungs, kidneys, and heart were aseptically dissected using sterile techniques. Specimens of lung, kidney, liver, and heart were fixed in 10% buffered formalin, paraffin-embedded, sectioned at 4 microns, and stained with haematoxylin and eosin (H&E). Sterile swabs were used for samples of lung, liver, spleen and kidney. Samples were inoculated into tryptone soy agar, blood agar (5% sheep blood enriched tryptone soy agar), and MacConkey agar, and incubated for 24–48 h at +37°C. Pure bacterial colonies of spherical or ovoid cocci, 1–2 μm in diameter, consistent with the genus *Photobacterium*, appeared on the blood agar plates at 48 hours post inoculation according Morick et al. (2023) (17). Bacterial species were confirmed using a rapid, proteomic based technique for identification of clinical bacterial isolates by protein profiling using matrix-assisted laser desorption ionization-time of flight mass spectrometry (MALDI-TOF MS) systems. Molecular characterization of bacterial species was conducted by PCR analysis. DNA was extracted from 1 gr of dissected lung, liver/spleen and kidneys using the CTAB method (9). PCR reactions were conducted to amplify regions of the 16S ribosomal DNA and *ureC* gene to assess the molecular attribution of *P. damselae* (17). To evaluate bacteria virulence, a PCR screening of haemolysin was conducted amplifying the pPHDD1 plasmid genes *damselysin* (*dly*) and *haemolysin A* (*hlyA*_*pl*_), and the chromosome gene *hlyA*_*ch*_ (1, 21). All PCR reactions were performed using the *DreamTaq* DNA polymerase (Invitrogen-ThermoFisher, Waltham, MA, USA) based on the manufacturer’s instructions. The nucleotide sequences of the primers used are listed in Table 1. Agarose gel (1.5%) electrophoresis was conducted on the amplification products at 100V and the GelDoc Go Imaging System (Bio-Rad, Hercules, CA, USA) was used for image acquisition. The amplification products of the 16S ribosomal DNA and *ureC* gene were directly sequenced by Sanger method (Eurofins Genomics, Ebersberg, Germany). PCR amplification using the primer pairs to amplify 16S, *ureC* and virulence genes produced fragments of the expected size (Figure 2, Table 1).

**Table 1:**
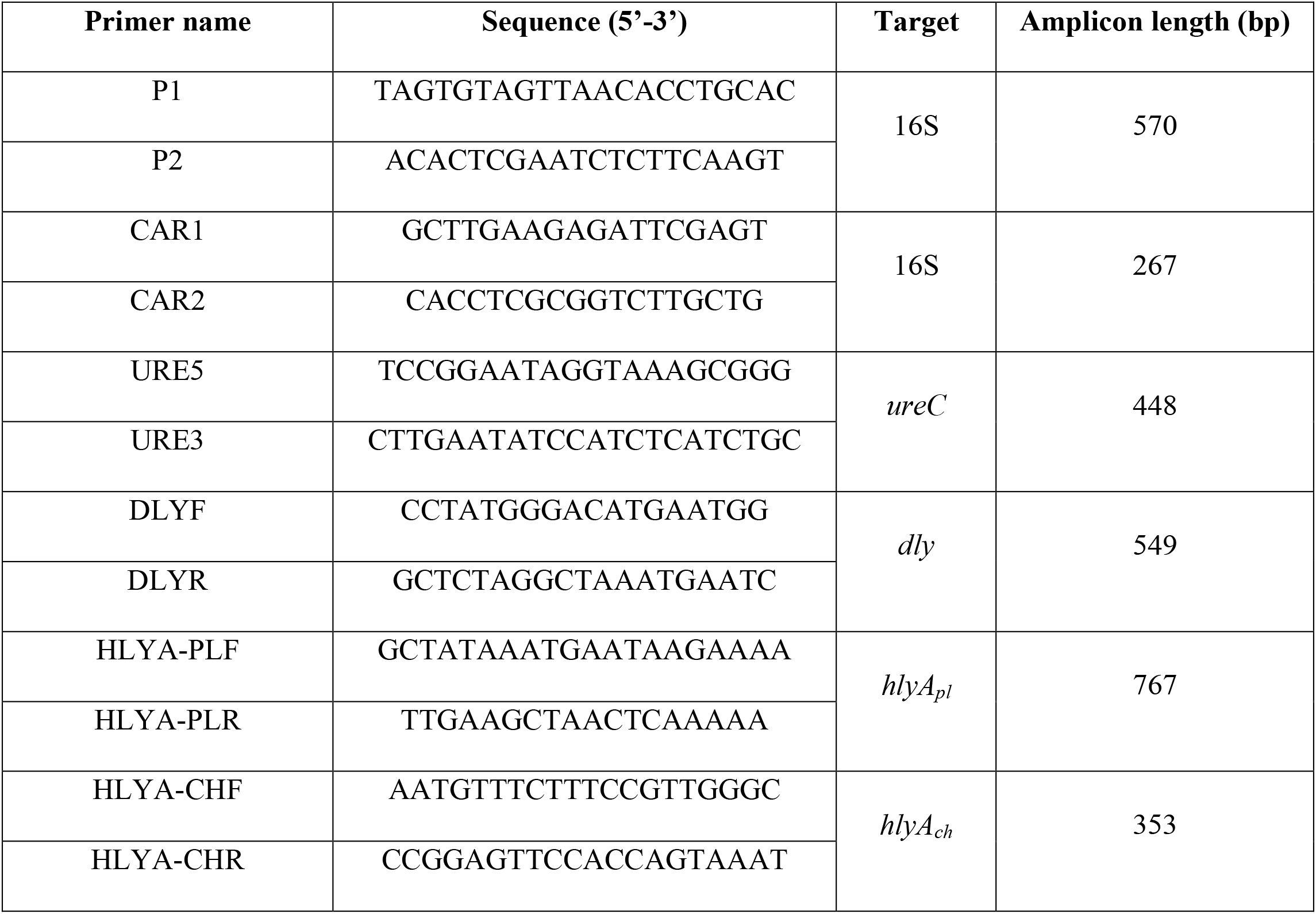
Nucleotide sequence of the primers used

**Figure 1.**
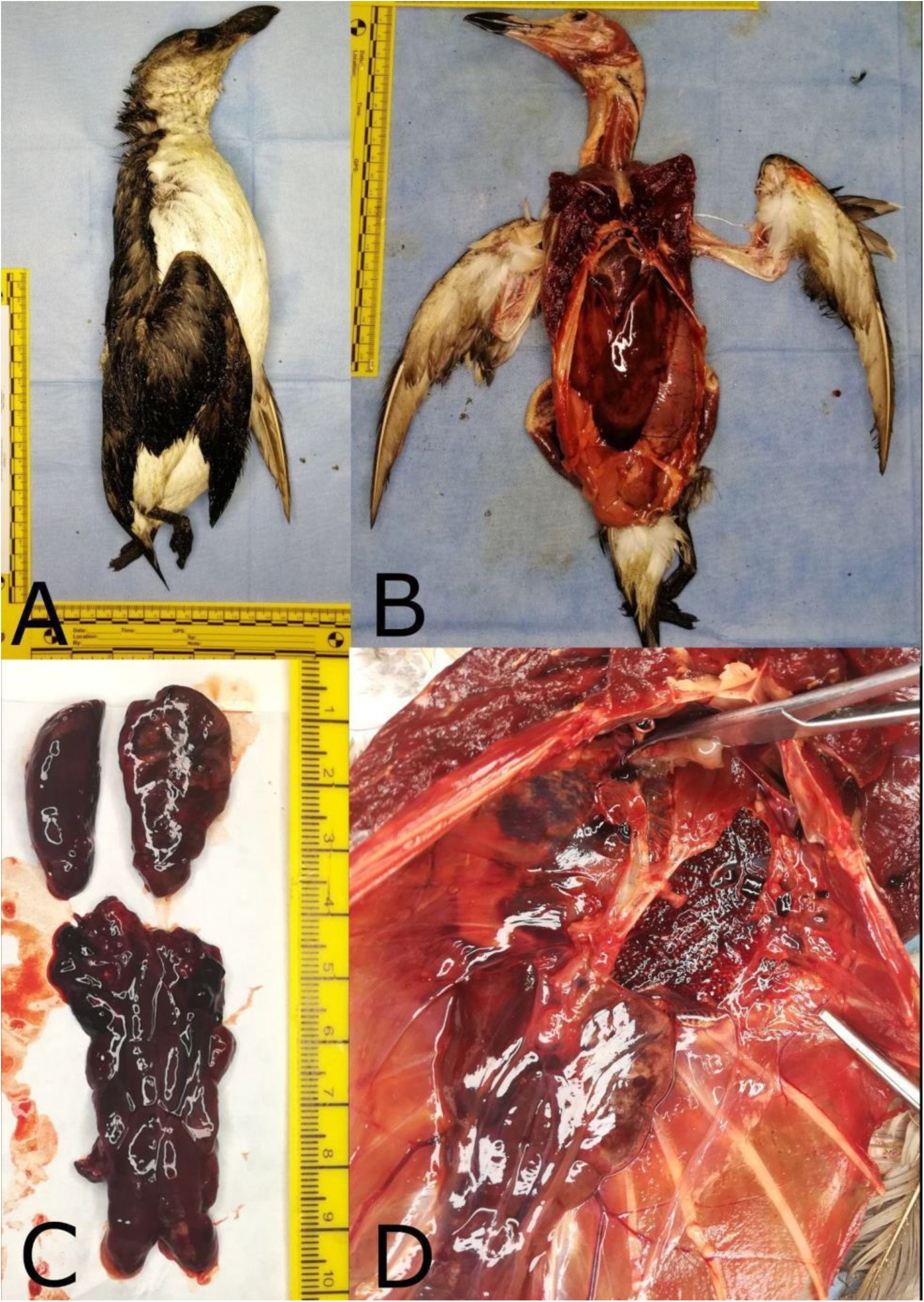
Gross pathologic examination of the Razorbill (*Alca Torda*, Linnaeus 1758) (A); Appearance of coelomitis (B) secondary to septicemia with severe congestion of liver, kidneys (C), and lungs (D).

**Figure 2:**
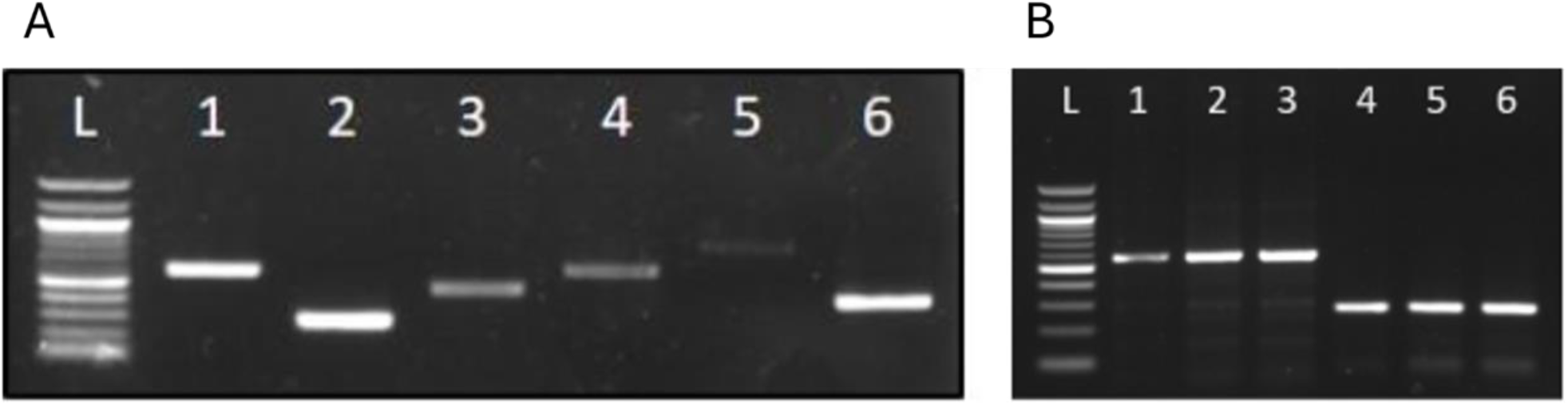
A: Agarose gel electrophoresis of the amplification products of the *Photobacterium damselae* genes from lung tissue. 100 bp DNA Ladder (NEB, Ipswich, MA, USA) (L); 16S (1, 2); *ureC* (3); *dly* (4); *hlyA*_*pl*_ (5); *hlyA*_*ch*_ (6). **B**: Agarose gel electrophoresis of the amplification products of *Photobacterium damselae* genes from liver, kidney and lung tissue. 100 bp DNA Ladder (NEB, Ipswich, MA, USA) (L); *16S* from liver (1, 4), kidney (2, 5) and lung tissue (lane 3, 6).

## Results

The post-mortem examination of the razorbill revealed coelomitis with severe congestion of the liver, lungs, kidneys, and myocardium. Bacterial isolation from lung, liver, and kidney samples showed the growth of pure spherical or ovoid cocci colonies on blood agar after 48 hours, consistent with the *Photobacterium* genus. Identification was confirmed via MALDI-TOF MS and further characterized molecularly using PCR, detecting the *ureC* gene and attributing the strain to *Photobacterium damselae* subsp. *damselae* (Pdd). Sequencing of the 16S gene fragments confirmed the presence of *P. damselae* in lung, liver, and kidney tissues. Additionally, PCR screening for virulence genes detected the *hlyAch* haemolysin gene in lung samples, suggesting a potential pathogenicity of the isolated strain. BLAST analysis of the sequenced 16S fragments confirmed the presence of *P. damselae* in the lung, liver/spleen and kidney tissue and the amplification of the *ureC* fragment permits its attribution to the *P. damselae* subsp. *damselae* strain. The obtained sequences were deposited in GenBank with the accession numbers PP946852 (16S), PP946853 (16S), and PP947936 (*ureC*). The amplification of the *haemolysin* genes shows that the Pdd strain found in the lung has virulence genes. Histopathologic examination of the liver showed a severe, multifocal perivascular inflammatory infiltrate predominantly composed of lymphocytes and plasma cells (Figure 3A). In the lung, the parenchyma was characterized by a mild inflammatory infiltrate, predominantly consisting of lymphocytes and plasma cells. Severe hemorrhages and blood vessels of irregular caliber, multifocally and severely filled with red blood cells (congestion), were also seen. A moderate amorphous and eosinophilic material (pulmonary edema) accumulation was observed within some alveolar spaces (Figure 3B). An inflammatory infiltrate in the kidney was predominantly observed, consisting of lymphocytes and plasma cells, coupled with numerous and widespread hemorrhages and tubular necrosis (Figure 3C). In the heart, the histological sections examined showed a mild and multifocal inflammatory infiltrate, primarily characterized by lymphocytes (Figure 3D).

**Figure 3.**
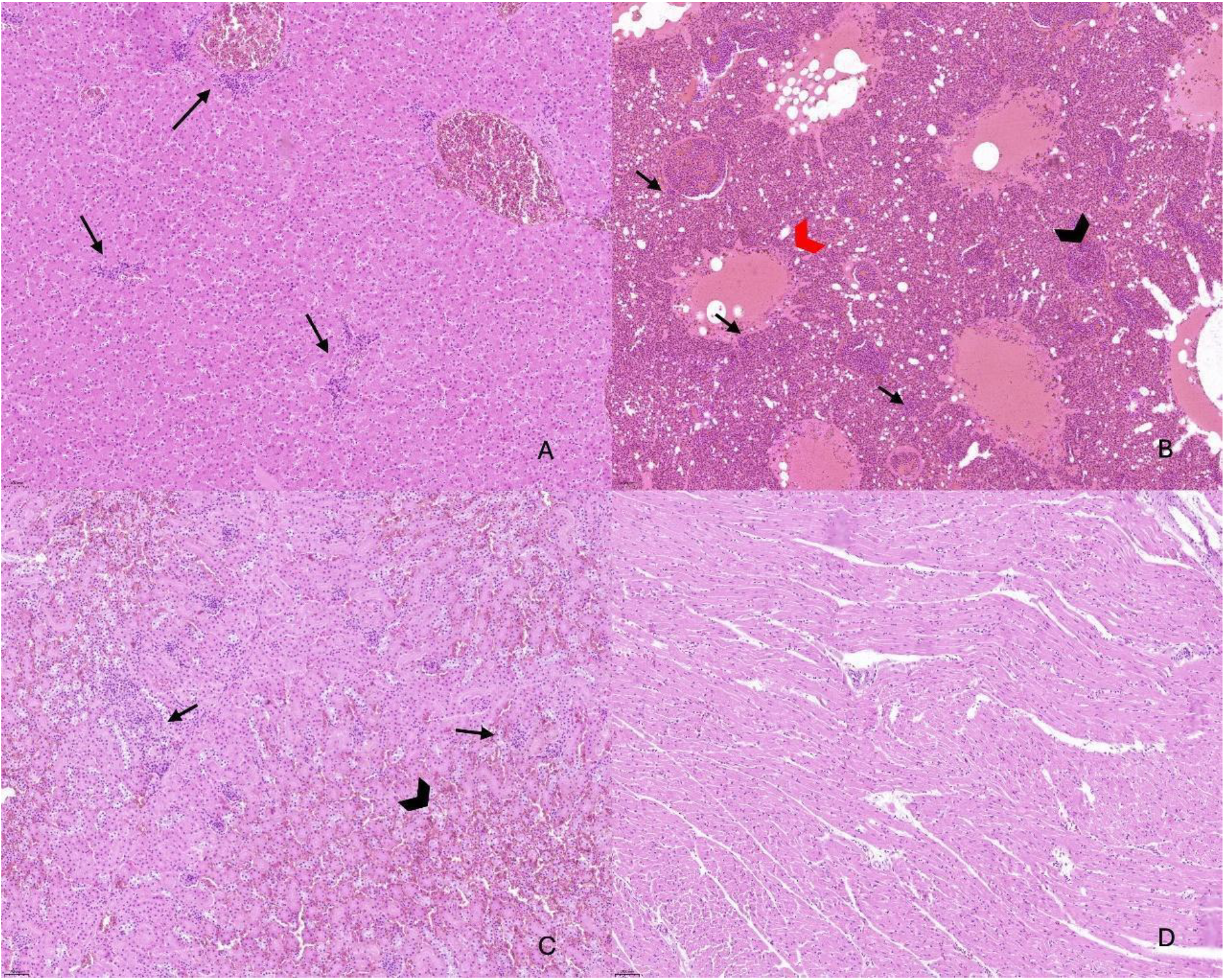
Histologic examination of the liver (A), lung (B), kidney (C), and heart (D) in Razorbill Bird. (A) The liver showed moderate, multifocal, and periportal infiltration of lymphocytes and plasma cells (arrows). (B) The lung showed capillary congestion (red arrowhead) and pulmonary edema (black arrowhead). Moderate infiltration of lymphocytes in the lung section was observed (arrows). (C) Numerous inflammatory cells (arrows) and diffuse hemorrhage (arrowhead) were observed in the kidney. (D) The heart showed few disseminated lymphocytes (arrows).

## Discussion

*Photobacterium damselae* subsp. *damselae* (Pdd) is commonly found in seawater, marine sediments, seaweeds, and marine organisms, thriving in warmer water temperatures ranging from 20 to 30 °C. This microorganism is recognized as a key pathogen affecting various species of both wild and farmed fish, leading to wound infections and hemorrhagic septicaemia. Pdd flourishes in coastal marine environments and has been extracted from seawater, seafood, seaweeds, and healthy marine animals (1, 19). However, this bacterium is primarily recognized for its ability to cause disease in various marine animals, including fish, mollusks, crustaceans, and cetaceans. (7, 22). It is nowadays considered an emerging pathogen in marine aquaculture, and numerous recent studies have reported its isolation from newly cultured species and from diverse geographical areas (13), where it is responsible for large economic losses. However, the prevention and control strategies put in place are still limited (11). Recently, this pathogen was reported to cause severe infection in humans causing opportunistic infections that may evolve into necrotizing fasciitis with fatal outcome. The majority of the reported infections in humans are caused by *Photobacterium damselae* subsp. *damselae*, and they typically originate from wounds exposed to salt or brackish water, usually occurring during the handling of fish or fishing tools (22). Unusual cases of infection have also been reported following the ingestion of raw seafood and through the urinary tract due to exposure to seawater. The majority of the cases occurred in coastal areas of the United States of America, Australia, and Japan (22). We documented the first isolation of *Photobacterium damselae* subsp. *damselae* in a razorbill but especially the first in a migratory bird. The finding of the razorbill specimen fits a context of irruption migration by seabirds observed along the coasts of the Mediterranean and the Tyrrhenian Seas during the winter of 2022\2023. This phenomenon has been attributed to extreme weather events in the North Atlantic, which are hypothesized to have displaced razorbills southward. Such displacement likely affected their physical condition, possibly due to difficulties in locating or catching prey. The status of these birds may have been further compromised by the suboptimal trophic characteristics of Mediterranean waters for this species (2). This may account for an increased susceptibility to infection by *Photobacterium damselae* subsp. *damselae*, as this microorganism is considered an opportunistic pathogen not only for humans, but also for a variety of marine animals, like cetaceans (4). It has also been demonstrated that warmer marine waters lead to the upregulation of virulence-associated gene transcription, thereby enhancing its pathogenic potential and increasing the likelihood of infection (15). Consequently, climate change is likely to impact not only the migratory and foraging patterns of razorbills and other seabirds (8, 18), but also to increase their exposure to potentially pathogenic agents such as *Photobacterium damselae* subsp. *damselae*. This isolation represents an important discovery as it highlights how this bacterium can infect a variety of marine animals and now also birds, expanding the ecological and health risk. The razorbill bird was found a few meters from the coast. Since these birds feed on fish, including those consumed by humans, and have been observed scavenging fishery waste at ports and requiring food directly from people (16), there is a potential risk of spreading this pathogen through seabirds, even over long distances. In fact, it is well established that migratory birds play a significant role in the ecology and transmission of zoonoses, potentially serving as both mechanical and biological vectors (10). This phenomenon poses a potential danger not only to marine fauna but also to fish farms and human health. Our results, together with those in the literature, highlight how this emerging pathogenic bacterium could cross species barriers with the risk of inducing disease in nearby and distant host populations with ecological and epidemiological implications for marine wildlife as well as human health. It also suggests that environmental changes, such as extreme weather events and warming waters, could facilitate the spread of emerging pathogens such as *Photobacterium damselae* subsp. *damselae*.

## Acknowledgments

The authors would like to thank the Local Health Authority (ASL) Napoli 1 Centro for their support in this study.

## References

1. Alba P, Caprioli A, Cocumelli C, Ianzano A, Donati V, Scholl F, Sorbara L, Terracciano G, Fichi G, Nocera FD, Franco A, and Battisti A. 2016. A new multilocus sequence typing scheme and its application for the characterization of Photobacterium damselae subsp. damselae associated with mortality in cetaceans. Front Microbiol 7:1656.

2. Balestrieri R, Vento R, Viviano A, Mori E, Gili C, and Monti F. 2023. Razorbills Alca torda in Italian Seas: A Massive Irruption of Historical Relevance and Role of Social Network Monitoring. Animals 13:656.

3. Barrett RT. 2015. The diet, growth and survival of Razorbill Alca torda chicks in the southern Barents Sea. Ornis Norvegica 38:25–31.

4. Battistini R, Masotti C, Giorda F, Grattarola C, Peletto S, Testori C, Zoppi S, Berio E, Crescio MI, Pussini N, Serracca L, and Casalone C. 2024. Photobacterium damselae subsp. damselae in Stranded Cetaceans: A 6-Year Monitoring of the Ligurian Sea in Italy. Animals 14:2825.

5. Boutabia L, Menaa M, Mederbal KE, Boulaouad BA, Ayyach K, Harzallah B, Missoum M, and Telailia S. 2023. Recent and exceptional irruption of the razorbill alca torda (linnaeus, 1758) on the algerian coastline. Nat Croat 32:233–239.

6. Carboneras C. 1988. The auks in the western Mediterranean. Ringing Migr 9:18–26.

7. Ceccarelli D, and Colwell RR. 2014. Vibrio ecology, pathogenesis, and evolution. Front Microbiol 5:256.

8. Diamond A, and Devlin C. 2003. Seabirds as indicators of changes in marine ecosystems: ecological monitoring on Machias Seal Island. Environ Monit Assess 88:153–175.

9. Doyle JJ, and Doyle LJ. 1999. Isolation of plant DNA from fresh tissue. Focus 13–15.

10. Georgopoulou I, and Tsiouris V. 2008. The potential role of migratory birds in the transmission of zoonoses. Vet Ital 44:671–677.

11. Gouife M, Chen S, Huang K, Nawaz M, Jin S, Ma R, Wang Y, Xue L, and Xie J. 2022. Photobacterium damselae subsp. damselae in mariculture. Aquacult Int 30:1453–1480.

12. Harrison P, Perrow MR, and Larsson H. 2021. Seabirds New Identification Guide. Bercelona, Lynx Editions, 0–600.

13. Labella AM, Arahal DR, and Castro D. 2017. Revisiting the genus Photobacterium: Taxonomy, ecology and pathogenesis. Int Microbiol 20:1–10.

14. Lavers JL, Jones IL, and Diamond AW. 2007. Natal and Breeding Dispersal of Razorbills (Alca torda) in Eastern North America.

15. Matanza X, and Osorio C. 2020. Exposure of the opportunistic marine pathogen Photobacterium damselae subsp. damselae to human body temperature is stressful condition that shapes the transcriptome, viability, cell morphology and virulence. Front Microbiol 11:1771.

16. Monti F, Mori E, Balestrieri R, Minichino A, Vento R, Viviano A, and Tiralongo F. 2024. First data on the diet of Razorbill Alca torda wintering in the Mediterranean Sea: Insights from social media. Mediterr Mar Sci 25:58–66.

17. Morick D, Blum SE, Davidovich N, Zemah-Shamir Z, Bigal E, Itay P, Rokney A, Nasie I, Feldman N, Flecker M, Roditi-Elasar M, Aharoni K, Zuriel Y, Wosnick N, Tchernov D, and Scheinin AP. 2023. Photobacterium damselae subspecies damselae Pneumonia in Dead, Stranded Bottlenose Dolphin, Eastern Mediterranean Sea. Emerg Infect Dis 29:179–183.

18. Orgeret F, Thiebault A, Kovacs KM, Lydersen C, Hindell MA, Thompson SA, Sydeman WJ, and Pistorius PA. 2022. Climate change impacts on seabirds and marine mammals: The importance of study duration, thermal tolerance and generation time. Ecol Lett 25:218–239.

19. Osorio CR, Vences A, Matanza XM, and Terceti MS. 2018. Photobacterium damselae subsp. Damselae, a generalist pathogen with unique virulence factors and high genetic diversity. J Bacteriol 200:e00002–18.

20. Pasquet E. 1988. Contribution à l’étude du régime alimentaire des guillemots de Troïl (Uria aalge) et petits pingouins (Alca torda) hivernant dans les eaux françaises. Alauda 56:8–21.

21. Rivas AJ, Balado M, Lemos ML, and Osorio CR. 2011. The Photobacterium damselae subsp. Damselae hemolysins damselysin and HlyA are encoded within a new virulence plasmid. Infect Immun 79:4617–4627.

22. Rivas AJ, Lemos ML, and Osorio CR. 2013. Photobacterium damselae subsp. Damselae, a bacterium pathogenic for marine animals and humans. Front Res Found.

